# Widespread parasite infections in living resident killer whales in the Northeast Pacific Ocean

**DOI:** 10.1101/2024.07.10.602791

**Authors:** Natalie C. Mastick, A.M. Van Cise, K.M. Parsons, E. Ashe, R. Williams, J.N. Childress, A. Nguyen, H. Fearnbach, J. Durban, C. Emmons, B. Hanson, D. Olsen, C.L. Wood

## Abstract

Multiple populations of resident ecotype killer whales (*Orcinus orca ater*) inhabit the Northeast Pacific, but the southern resident killer whale (SRKW) population is the most at-risk. SRKWs were listed as endangered in the United States in 2005 and have since shown little sign of recovery. Several factors have been identified as key threats to this population, and previously published studies suggest the population may be energetically stressed. Underlying health risks, such as parasitism, may be contributing to this population’s failure to recover, but little is known about parasite infections in living individuals from natural killer whale populations. To assess the prevalence of internal parasite infections in Northeastern Pacific killer whales, we examined scat from endangered SRKW (n = 25) compared to two conspecific populations of resident killer whales that are not in decline: northern resident (NRKW, n = 2) and southern Alaska resident killer whales (SARKW, n = 7), and one offshore killer whale (OKW, n = 1). We analyzed 35 fecal samples collected from 27 wild killer whales using both microscopic identification of parasite eggs and genetic detection of parasites through DNA metabarcoding. We used body condition indices derived from concurrent aerial photogrammetry to evaluate whether parasite infection status was associated with individual body condition. We found that most individuals sampled (94%) were positive for *Anisakis* spp. – a parasitic nematode known to inhabit the intestines of cetaceans. These infections were detected across populations, and were not correlated with body condition, based on limited paired data. These results suggest that *Anisakis* infection is widespread among resident killer whales of the Northeast Pacific. The widespread detections of Anisakis among the samples examined here emphasizes the need for further work to understand the potential health impacts of parasitic infections on individual killer whales, and potential synergistic effects with other environmental stressors.

## INTRODUCTION

Among sublethal stressors, both internal and external parasites are known to impact the health of wildlife populations (Hudson et al. 1998; Tompkins and Begon 1999). Parasites can work in tandem with other stressors to reduce the health of their hosts; parasites both reduce the energy available to hosts (Shanebeck et al. 2022) and modulate the immune response either indirectly by diverting energy away from the immune system or directly by manipulating the immune system (Schmid-Hempel 2008). Together, these impacts can result in a host that is more vulnerable to other infections, or more vulnerable to the impact of other stressors (Beldomenico et al. 2008; Marcogliese and Pietrock 2011). This can lead to a negative feedback loop, in which hosts are unable to resist infection after being subjected to stress from multiple factors, which reduces their energetic condition and leaves them more vulnerable to stress (Beldomenico and Begon 2010). For example, a study on harbor porpoises (*Phocena phocoena*) found that nutritionally stressed individuals were more likely to have parasite infections (Ten Doeschante et al. 2017). In this study, the authors suggested that parasites may have a greater impact on fitness in unhealthy animals (Ten Doeschate et al. 2017). Additionally, host characteristics such as age or sex may influence susceptibility to parasite infections where parasitism disproportionately affects some members of the population (Marcogliese and Pietrock 2011). Parasitism can lead to significant and long-lasting energy loss in threatened or stressed populations (Shanebeck et al. 2022), which can subsequently negatively impact reproductive success and population growth (Irvine 2006; Riordan et al. 2007).

One of the most common parasites found in the intestinal tract of marine mammals at necropsy are nematodes (i.e., roundworms) in the family Anisakidae (hereafter anisakids; see Table 1 for a list of known delphinid parasites; Dailey 2001). There are three prominent genera that use marine mammals as their definitive hosts: *Anisakis* spp., which infect cetaceans, *Pseudoterranova* spp., which infect pinnipeds, and *Contracaecum* spp., which infect pinnipeds and seabirds (Køie et al. 1995; Klimpel and Palm 2011). Thirty-four species of cetaceans are known to harbor *Anisakis* spp., including killer whales (Mattiucci and Nascetti 2007; Klimpel and Palm 2011; Raverty et al. 2020). The *Anisakis* spp. life cycle involves multiple larval phases and takes place mainly in the pelagic environment (Klimpel and Palm 2011; Figure 1). Eggs are deposited into the ocean through cetacean feces, where they develop into larvae and are transported up the food web via ingestion, infecting crustacean and fish intermediate and paratenic hosts (Klimpel and Palm 2011). Cetaceans become infected by ingesting infected intermediate hosts, at which time the larvae develop into adults and reproduce within the definitive host’s gastrointestinal tract.

**Figure 1:**
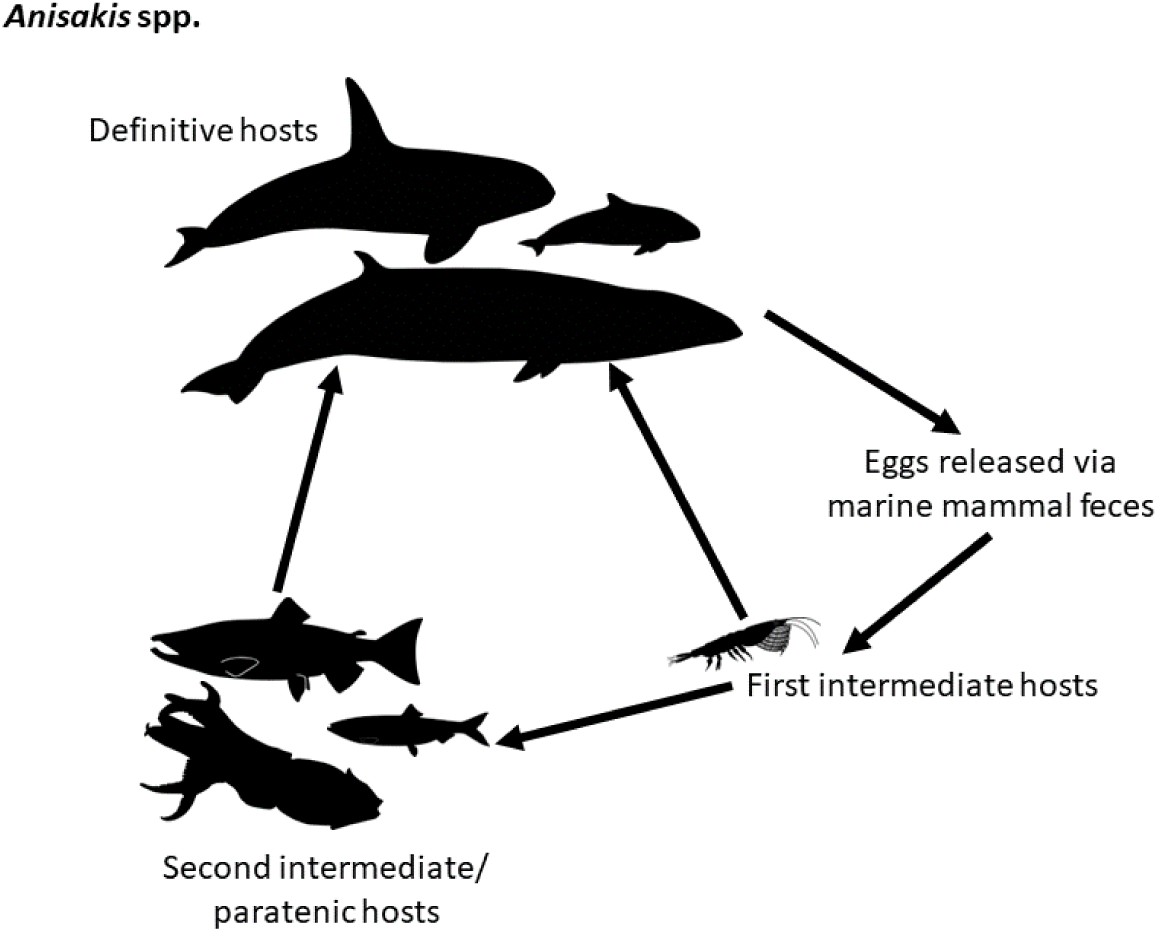
Life cycle of *Anisakis* spp. *Anisakis* spp. infect cetaceans as their definitive hosts, and after their eggs develop in the water column, larvae are consumed by an invertebrate which serves as their first intermediate host. The larva is transmitted to paratenic hosts when consumed by a fish or cephalopod - hosts in which the larvae does not develop, but which can help the parasite get to the definitive host. All images obtained from PhyloPic. Vector images of cetaceans courtesy of Chris Huh under the Creative Commons Attribution-ShareAlike 3.0 Unported license.

**Table 1:**
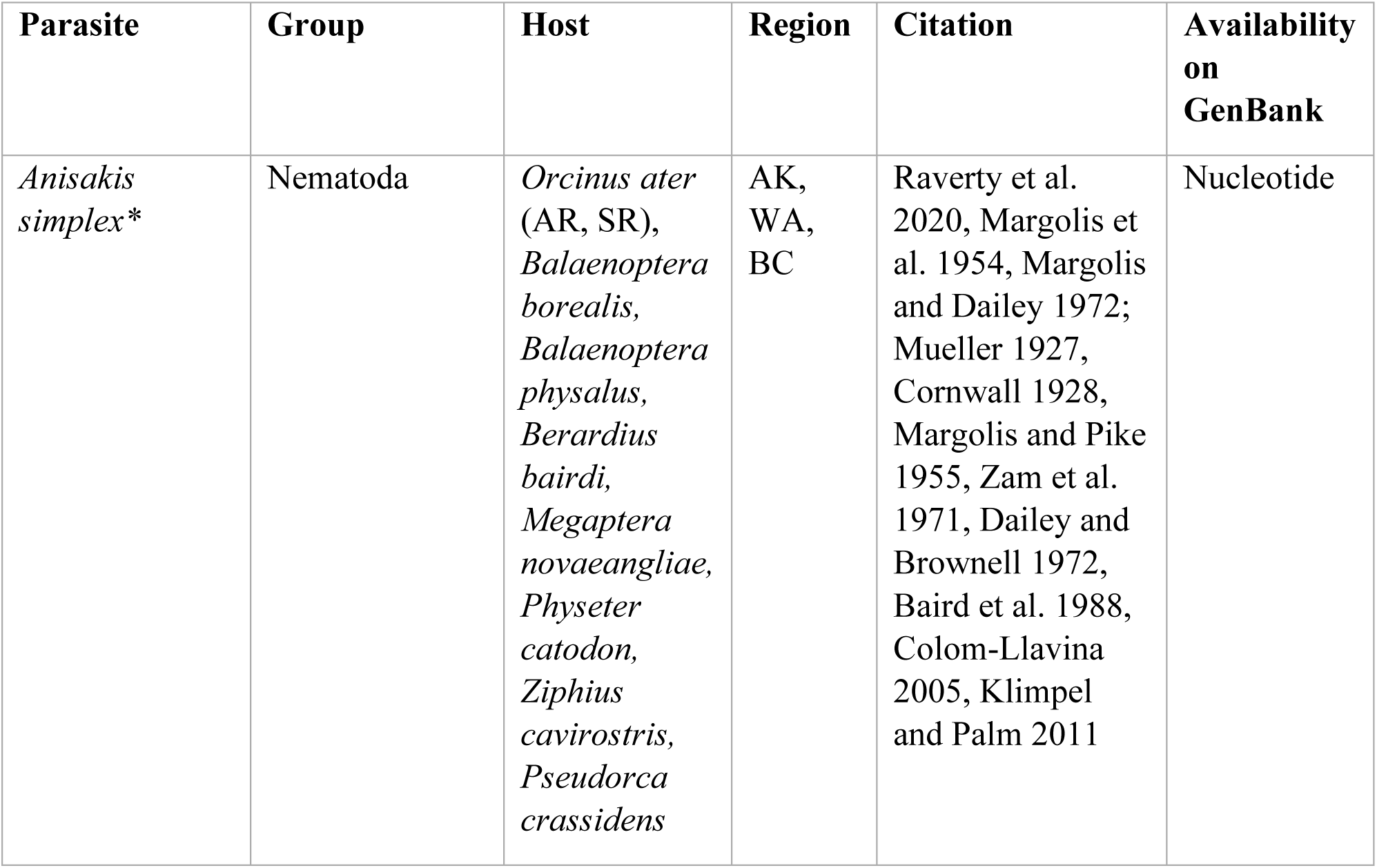
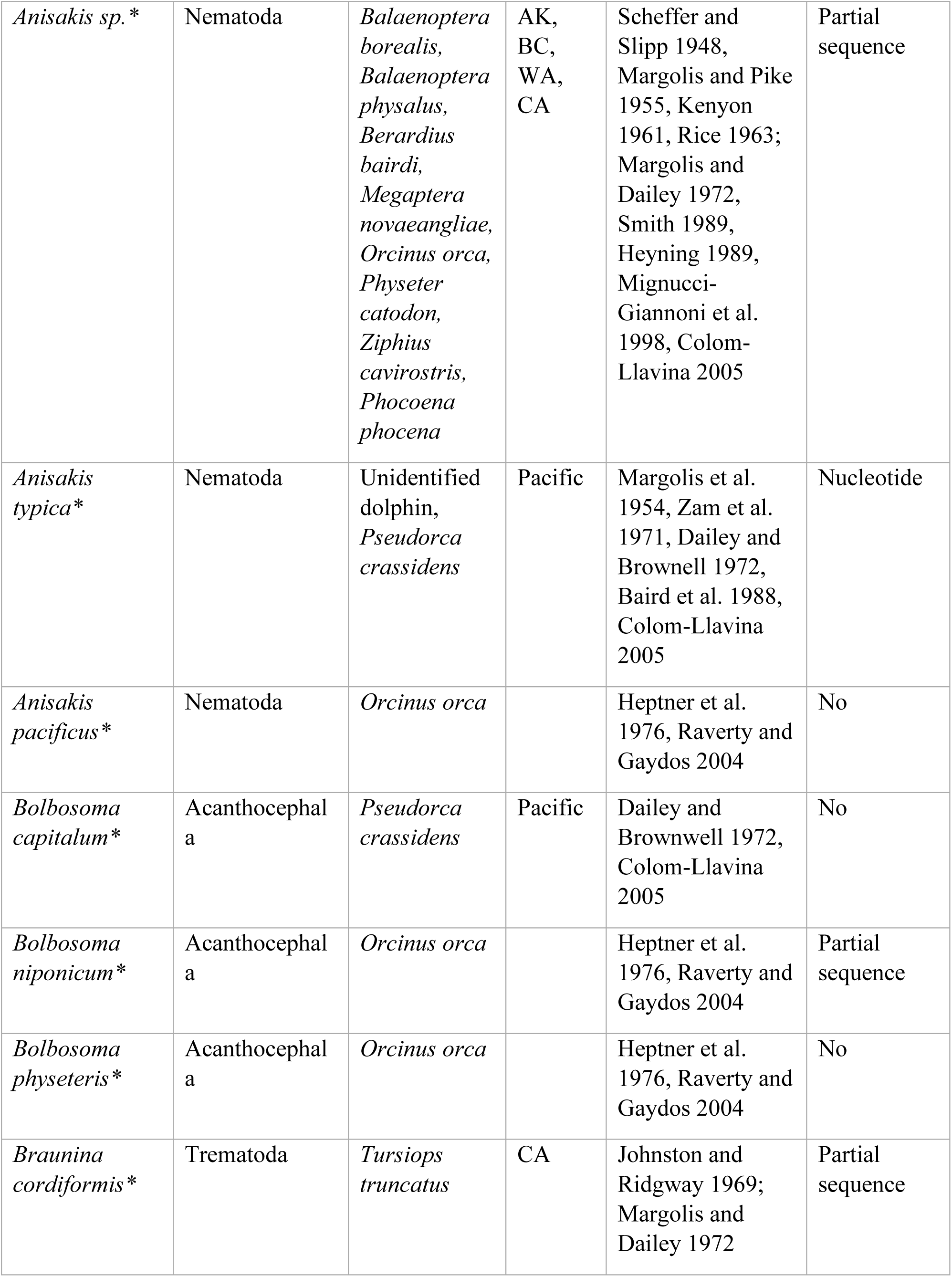
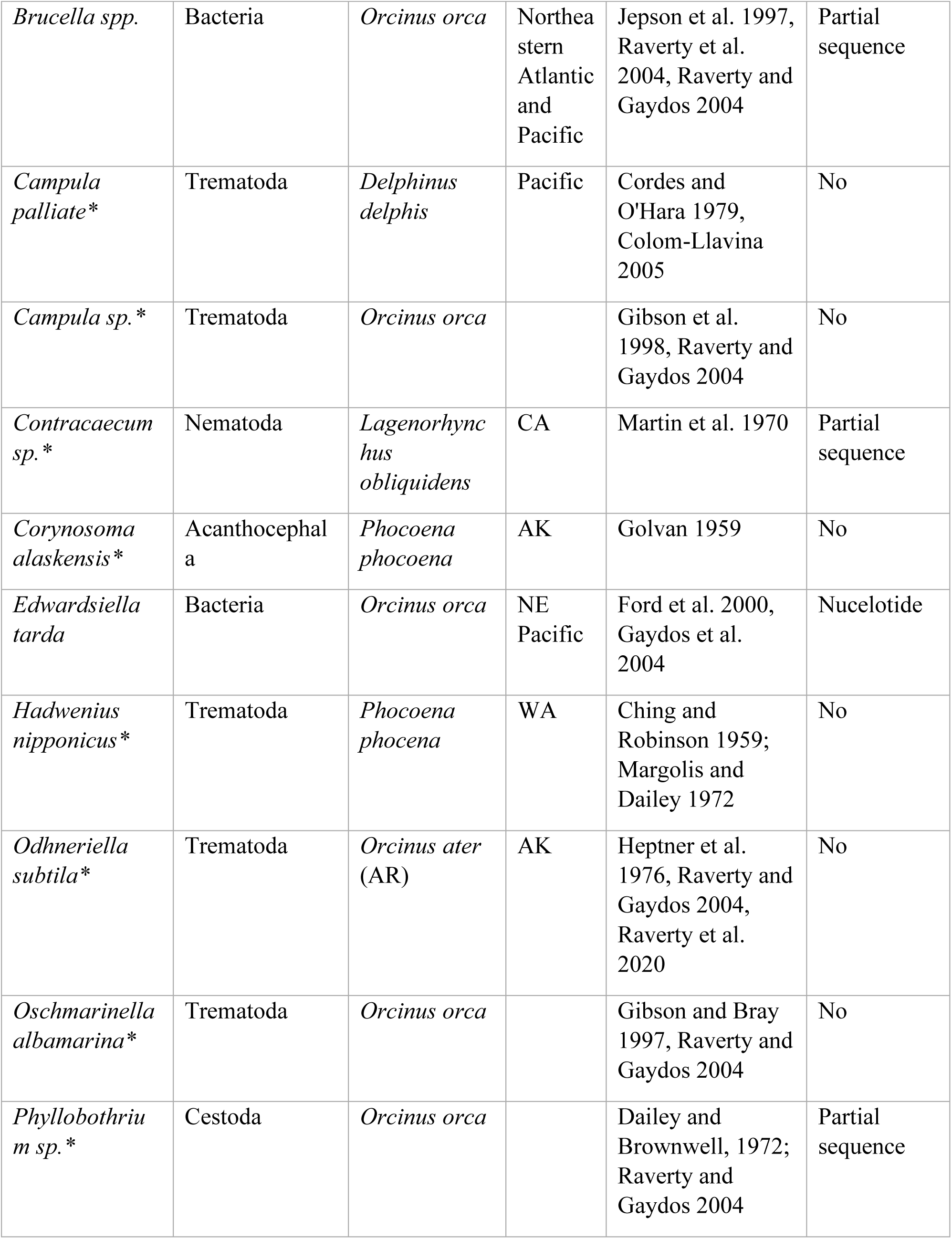
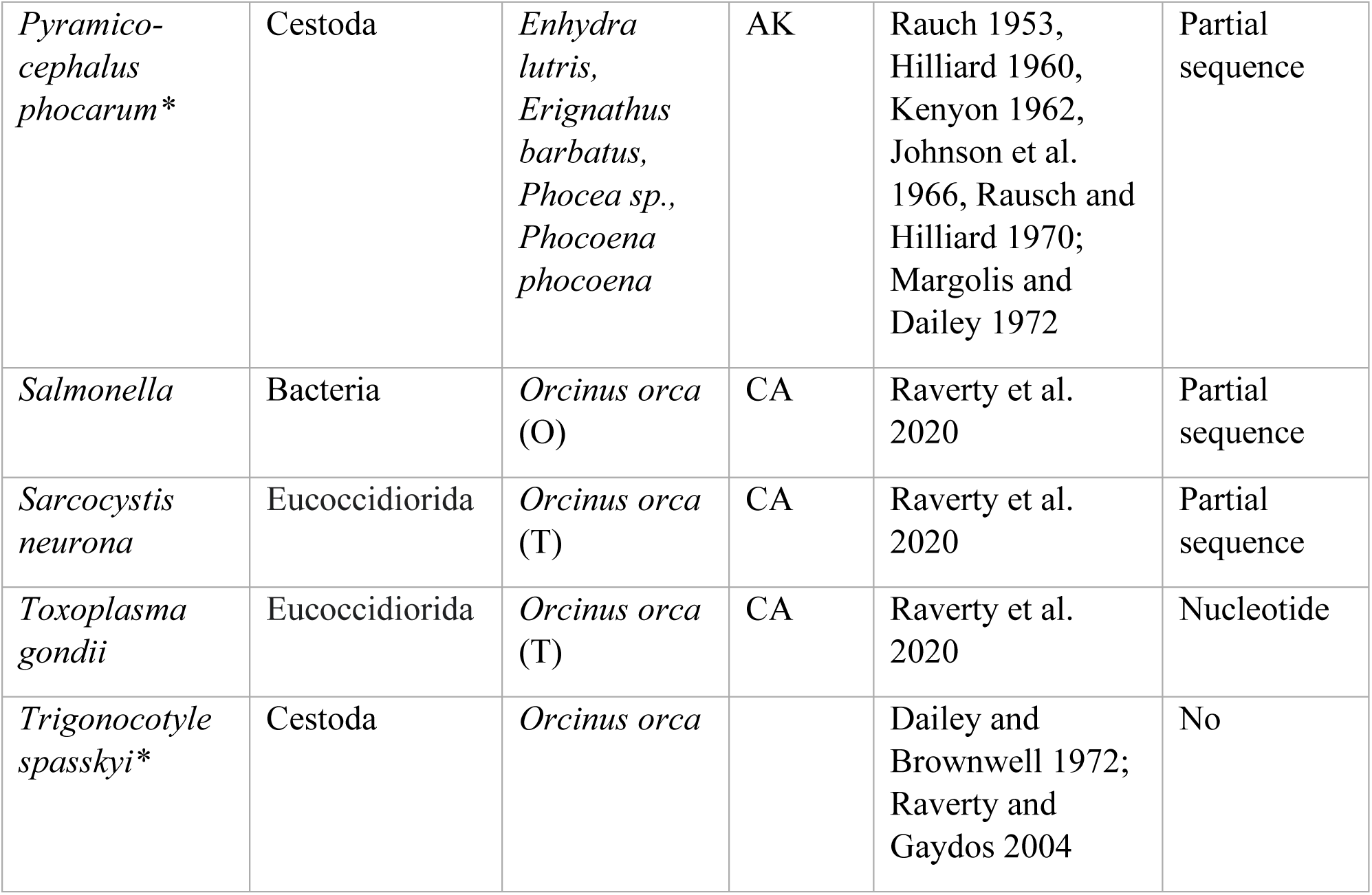
List of parasites that infect Delphinidae in the Northeast Pacific, compiled from published necropsy reports. Parasites that were likely to infect the gastrointestinal tract are indicated with an asterisk.

Anisakids cause both direct and indirect fitness costs in their marine mammal definitive hosts. *Anisakis* spp. can cause gastritis and ulceration (Cattan et al. 1976; Haebler & Moeller 2021), and they can cause peritonitis that ultimately leads to hemorrhaging and death (Dailey and Stroud 1978; Stroud and Roffe 1979; van Beurden et al. 2015). Though they rarely cause mortality, anisakids probably have an underestimated effect on host health as an energy sink (Shanebeck et al. 2022).

A recent study showed that there has been an increase in *Anisakis* spp. abundance in prey of marine mammals globally (Mastick et al. 2024a). In Puget Sound, abundance of the anisakid *Contracaecum* spp. in fish has increased with increasing marine mammal abundance (Mastick et al. *in revision*). This presents the opportunity for rising rates of anisakid infections, which may pose a threat to many marine mammal species, but those whose populations are already declining because of cumulative stressors could be at particularly high risk.

There are several populations of resident killer whales (*Orcinus orca ater*, hereafter RKWs) in the Northeast Pacific Ocean, including southern Alaska (SARKWs), northern (NRKWs), and southern resident killer whales (SRKWs). These populations have partially overlapping ranges that span the west coast of the United States and Canada, from the Gulf of Alaska to California, and share piscivorous diets consisting primarily of salmonids during the summer months (*Oncorhynchus* spp.; Bigg 1982; Krahn et al. 2004; Matkin 2011; Ford and Ellis 2006; Hanson et al. 2021). While they overlap in range and diet, SARKW and NRKW populations have increased in abundance while SRKWs have declined (Olesiuk et al. 2005; Fisheries and Oceans Canada 2018; Murray et al. 2021; Matkin et al. 2014; Lacy et al. 2017).

Declines in the abundance of SRKWs are thought to be attributable to inbreeding (Kardos et al. 2023) and the cumulative effects of multiple stressors (Ford et al. 2010; DFO 2017; NMFS 2008; Nelson et al. 2024). SRKWs, and killer whales in general, have a relatively low intrinsic population growth rate, which leaves them particularly susceptible to stress (Stark et al. 2004).

While an individual might tolerate the impacts of one sublethal stressor, the synergistic effects of multiple stressors can compound one another, leading to greatly diminished fitness and fecundity (Williams et al. 2016; Wright, 2012). Lacy et al. (2017) modeled the cumulative effects of known stressors of SRKWs and predicted a stable population — a prediction that contrasts with the declines observed since 2016 (Marine Mammal Commission 2023). However, the applied model was limited to recognize extrinsic stressors and did not capture additional sublethal stressors and intrinsic factors affecting SRKWs. If SRKWs within a specific age class or sex were particularly susceptible to parasitism, then parasitism could explain the failure of SRKWs to maintain the stable population predicted by Lacy et al. (2017).

Among wild cetaceans, the prevalence and impact of parasitism as a sublethal stressor is unknown due to the difficulty of assessing parasite infections in wild, living animals. This is further exacerbated by the challenges associated with collecting fecal samples from free-swimming cetaceans. Because of these challenges, modern parasitological examinations are limited to analyzing a limited number of fecal samples from wild animals or conducting necropsies of deceased animals, the latter likely being unrepresentative of healthy wild individuals due to biases associated with cause of death (Dailey and Stroud,1978; Aguilar and Borrell 1994; Ten Doeschate et al. 2017; Hermosilla et al. 2018; Raverty et al. 2020). Through necropsies, we know that marine mammal carcasses can carry high loads of parasites (Dailey and Stroud 1978; Dailey 1980; Stroud and Roffe 1979), but the parasite burden of living cetaceans is largely unknown (e.g., Raverty et al. 2017; Raverty et al. 2020; Lehnert et al. 2023).

Here, we present data on the parasite infections found in resident killer whales based on the analysis of fecal samples collected from living killer whales in the Northeast Pacific Ocean, with a focus on endangered SRKWs. To assess the extent and potential impact of parasite infections, we examined data from body condition assessments and fecal samples to characterize the types of parasites that infect RKWs, compare the types and prevalence of parasitic infections among RKW populations in the region, and test whether body condition is associated with the presence of parasites in SRKWs. Our findings constitute the first assessment of parasites in living cetaceans in the Northeast Pacific.

## METHODS

### Fecal Sample Collection

Resident killer whales in the Northeast Pacific have been photo-identified and monitored since the early 1970s, making them one of the most well-studied cetaceans (e.g., Center for Whale Research, North Gulf Oceanic Society, Fisheries and Oceans Canada’s Pacific Biological Station). Routine monitoring and health assessments of SRKWs have allowed for the collection of long-term health data in this population, including fecal samples, dietary preferences, and, for SRKWs, body condition estimates derived from photogrammetry data (Fearnbach et al. 2020, 2018; Ford et al. 2016; Hanson et al. 2010). Fecal samples were collected from killer whales following a standardized protocol during routine fieldwork led by teams conducting long-term health assessments of SRKWs (Supplementary Materials 1). After genotyping to assign fecal samples to individual SRKWs (Ford et al. 2018), 1-mL fecal subsamples were aliquoted from SRKW (n = 25), SARKW (n = 7), NRKW (n =2) samples, and from one offshore killer whale (OKW) sample for intestinal parasite analyses. SARKW were genetically identified and known to be unique individuals. NRKW and OKW samples are from unknown individuals, but were assumed to be unique individuals. All samples were associated with a specific collection date and location.

### Creating a parasite identification guide for North Pacific wild killer whales

To determine which parasites may infect killer whales, we reviewed the literature for documentation of intestinal parasites that infect members of the family Delphinidae in the Northeast Pacific Ocean, which resulted in our list of probable species (Table 1). This list was used to assemble an identification guide, including photos or written descriptions of parasite eggs (Supplementary Materials 2). This list was also used to query GenBank® (www.ncbi.nlm.nih.gov/genbank/) to identify species with publicly accessible genetic sequence data (Table 1) and identify target loci for DNA metabarcoding analysis.

### Fecal float and sedimentation analysis

For each sample, we performed both fecal floatation and sedimentation. Each fecal sample was partially thawed, and a subsample was collected with a flame-sterilized metal spoon and weighed. We aimed to take 0.5-g samples (wet weight), but if the fecal sample was less than 1 g, we took approximately half of the sample and recorded the exact mass. We subjected each subsample to standard fecal sedimentation and floatation protocols (Girard et al. 2016, Supplementary Materials 1). Two trained observers later identified the eggs using a published marine mammal parasite egg identification key (Dailey et al. 1980) – and when available, images of parasite eggs that infect other marine mammals in the northeast Pacific (Supplementary Materials 2). To prepare the data for downstream modeling, we classified individual infection status as a binomial response — uninfected or infected — reflecting whether eggs were present in a sample.

### Fecal DNA metabarcoding analysis

Additionally, we ran an independent molecular genetic approach to detect parasites and pathogens. We amplified and sequenced a 156 bp sequence of the 28S rRNA using previously published primers (nucLSUDf1 and nucLSUDr1; Sonnenberg et al. 2007; Cabodevilla et al.

2022); this region includes species-specific variability that allows detection and classification of most marine and terrestrial helminths. DNA extraction, library preparation, sequence metabarcoding, and bioinformatic processing were conducted by Jonah Ventures (Boulder, CO) using a small portion of each fecal sample. For detailed information on the laboratory protocols and bioinformatic pipelines used to generate metabarcoding data see Supplementary Materials 1.

### Photogrammetry data

Aerial images of SRKWs were collected by an aerial observer from a helicopter in 2013 and using a remotely operated APH-22 hexacopter drone (Aerial Imaging Solutions, Old Lyme, CT) with an affixed Olympus E-PM2 camera between 2015–2021 (Fearnbach et al. 2011, 2018, 2020; Durban et al. 2015). Photogrammetry data were assigned to individual whales using an established aerial catalog showing markings that are visible from the air, allowing measurements to be linked to whales of known age and sex (Fearnbach et al. 2011, 2020; Durban et al. 2015). The eye patch ratio, a ratio of two different length measurements taken between the inside edges of the white eye patches at 75% of their length compared to between their anterior edges on the head of a killer whale, was used as a quantitative metric of body condition (Figure 2, e.g., Fearnbach et al. 2020). Body condition was classified into one of five categories based on measurements, as described by Stewart et al. (2021). To link parasite infection status with body condition, we compiled body condition data collected within a month of each scat sample. This resulted in 19 fecal-to-body-condition pairs for 14 unique whales.

**Figure 2:**
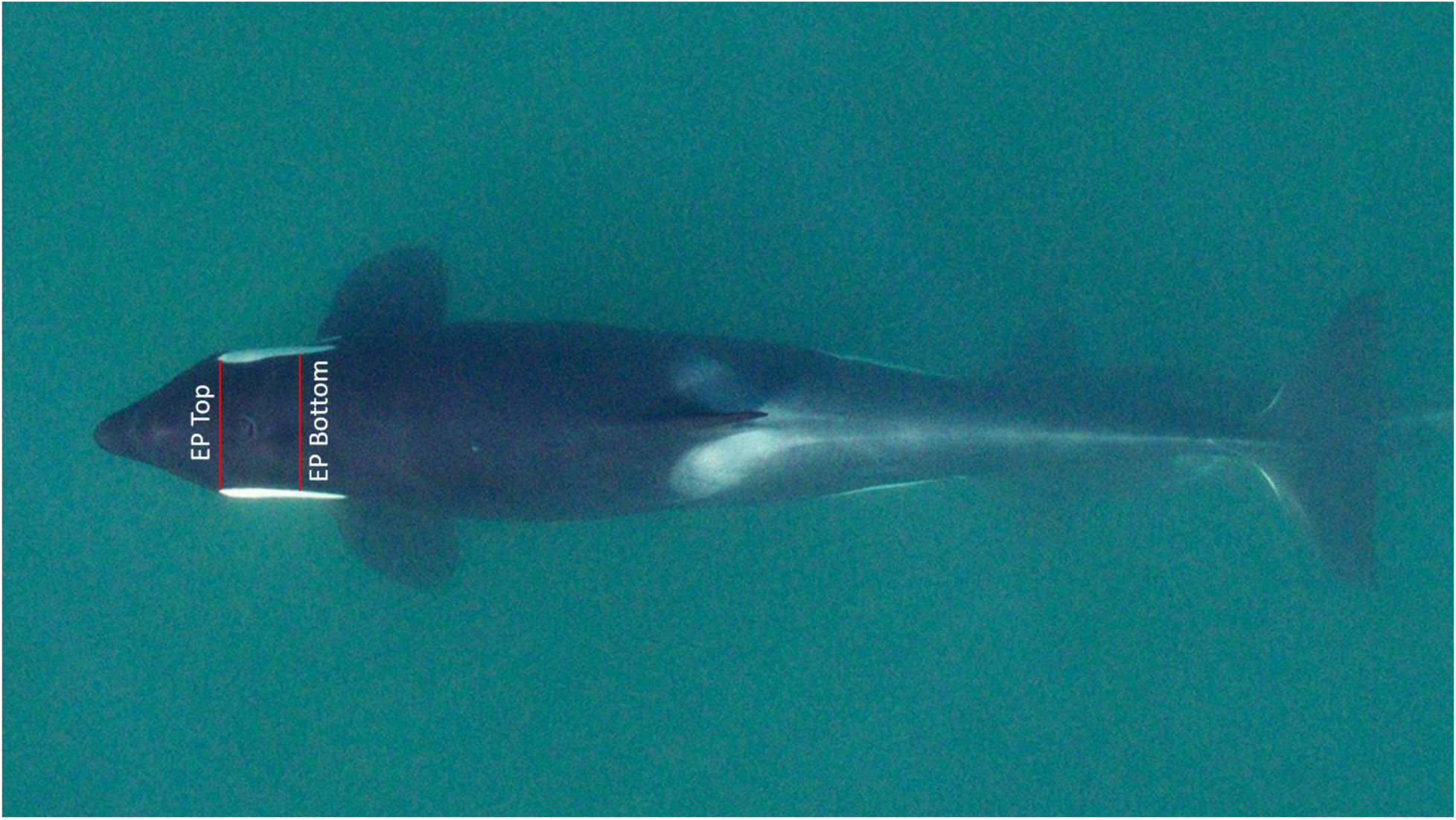
Aerial image showing the measurements for the eye patch ratio (EPR, proportion of “EP Top” and “EP Bottom”). The EPR is an indicator of nutritional condition, allowing changes in condition to be detected on both a seasonal and annual level.

### Statistical analysis

We had three objectives in our statistical analysis: 1) Determine whether parasite infection status differed between two of the three resident killer whale populations; 2) determine whether parasite infection status was correlated with temporal (i.e., month or year) or demographic (i.e., age, sex, or pod) factors within the SRKW population; and 3) identify whether body condition varied with parasite infection status within the SRKW population. The final two objectives were only assessed using data and samples from the SRKW population due to the low number of samples from the other two RKW populations, which precluded meaningful statistical analysis. For each model, we performed model selection by comparing the full model to models with every other combination of fixed effects using the MuMIn package in R (Bartoń 2023) and selected the model with the lowest AIC score.

As egg presence/absence is a regularly used metric for assessing parasite infection (Ten Doeschate et al. 2017), we used morphological presence/absence as the response variable in our models. We ran our models on parasite infection status using morphological data, because the ratio of infected to uninfected whales diagnosed through genetic data was too great for the models to successfully converge. To determine whether infection status differed by population, we ran a generalized linear mixed-effect model (GLMM) for each parasite genus detected in multiple whales through morphological identification using the glmmTMB() function in the R package of the same name (Brooks et al. 2017). Model 1 included infection status as the response variable, population (SRKW, SARKW, or NRKW) as a fixed effect, and year and individual identity as random effects. We included most available morphological data (*n* = 34); we excluded the sample collected from OKW, which provided too small of a sample size (*n* = 1) to include in the model.

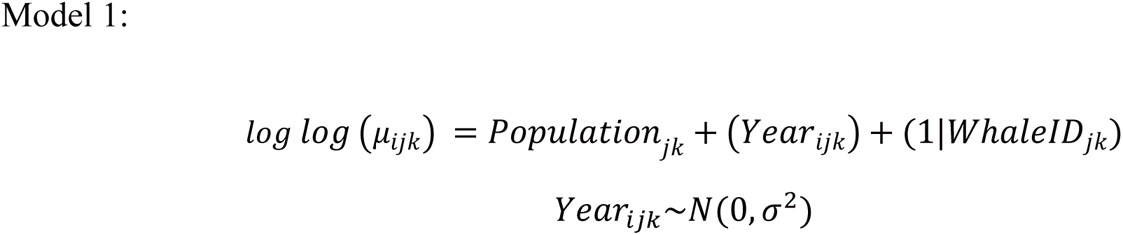

Where the response variable 𝜇_𝑖𝑗𝑘_ represents presence or absence of a parasite from the *i*th sample from the *j*th whale in the *k*th year when the sample was collected.

To determine if parasite infection status was affected by demographic or temporal factors, we ran a GLMM for each parasite taxon using SRKW data excluding whales from K pod, as there were too few whales sampled from K pod (N = 1 sample from K pod), resulting in 23 samples. We incorporated both demographic and temporal variables as fixed effects in the full model: age class (juvenile, subadult, or adult), sex, pod (J or L pod only), month, and year. Year was scaled using the *scale()* function in base R prior to model fitting. Whale ID was included as a normally-distributed random effect.

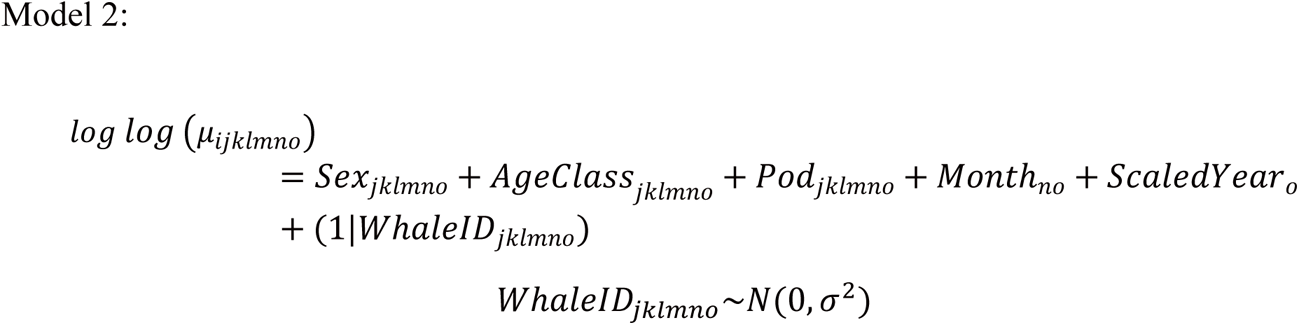

Where the response variable 𝜇_𝑖𝑗𝑘𝑙𝑚𝑛𝑜_ represents a binary indicator of presence or absence of a parasite from the *i*th sample from the *j*th whale of the *k*th sex and the *l*th age class in the *m*th pod sampled in the *n*th month of the *o*th year.

To determine if infection status varied by body condition, we restricted the dataset to SRKW with a corresponding body condition metric (*n* = 19). Our response variable was the infection status for each parasite. We ran a GLMM with a binomial distribution using the glmmTMB package. Fixed effects were body condition class (a categorical class between 1 and 5), pod, sex, and age class. Whale ID was included as a normally-distributed random effect.

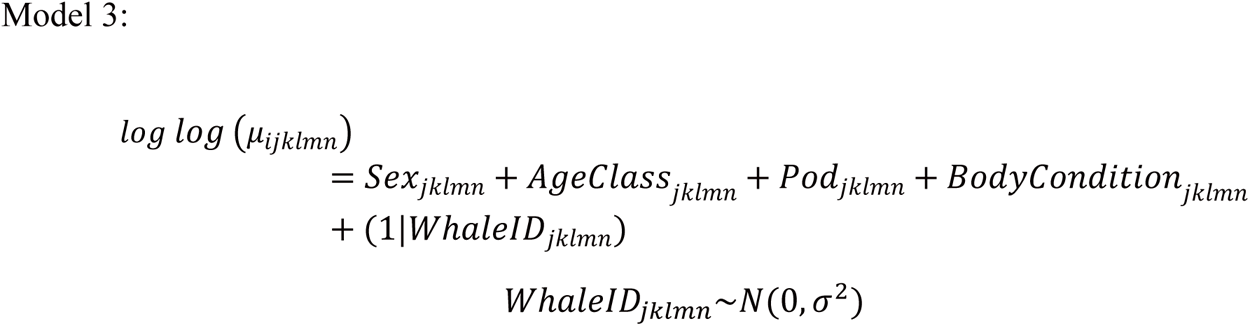

Where the response variable 𝜇_𝑖𝑗𝑘𝑙𝑚𝑛_ represents the relative abundance of each parasite from the *i*th sample collected from the *j*th whale of the *k*th sex in the *l*th pod with the *n*th body condition class.

To assess if parasite infection status had a significant effect on body condition, we also ran an ordinal regression model with body condition class as the response variable, parasite infection status, pod, and sex as fixed effects, and age class and whale identity as random effects using the clmm() function in the ordinal package in R (Christensen 2022). Finally, we ran a power analysis to measure the statistical power of Model 3 to detect a correlation if one existed (Supplementary Materials 1).

## RESULTS

### Fecal floatation and sedimentation analysis

Nematodes in the Anisakidae family were the only parasites found in both fecal floatation and sedimentation, with 2,732 eggs identified (in 26/35 killer whale samples). Based on the diagnostic key (Dailey et al. 1980), we identified two genera in fecal floatation and sedimentation analysis: *Contracaecum spp.* (*n*_SARKW_ = 23, *n*_NRKW_ = 0, *n*_SRKW_ = 228) and *Anisakis* spp. eggs (*n*_SARKW_ = 105, *n*_NRKW_ = 1, *n*_SRKW_ = 2375; Figure 3). We grouped eggs identified as *Anisakis* spp. and *Contracaecum* spp. into the family Anisakidae to report a single combined morphological egg count per float and a single combined egg count per sedimentation. Generally, we detected more anisakid eggs in killer whale fecal samples using fecal sedimentation than fecal floatation. There was only one instance in which the number of anisakid eggs identified in the fecal floatation exceeded the number identified in the fecal sedimentation (1 egg detected in the floatation, 0 in the sedimentation).

**Figure 3:**
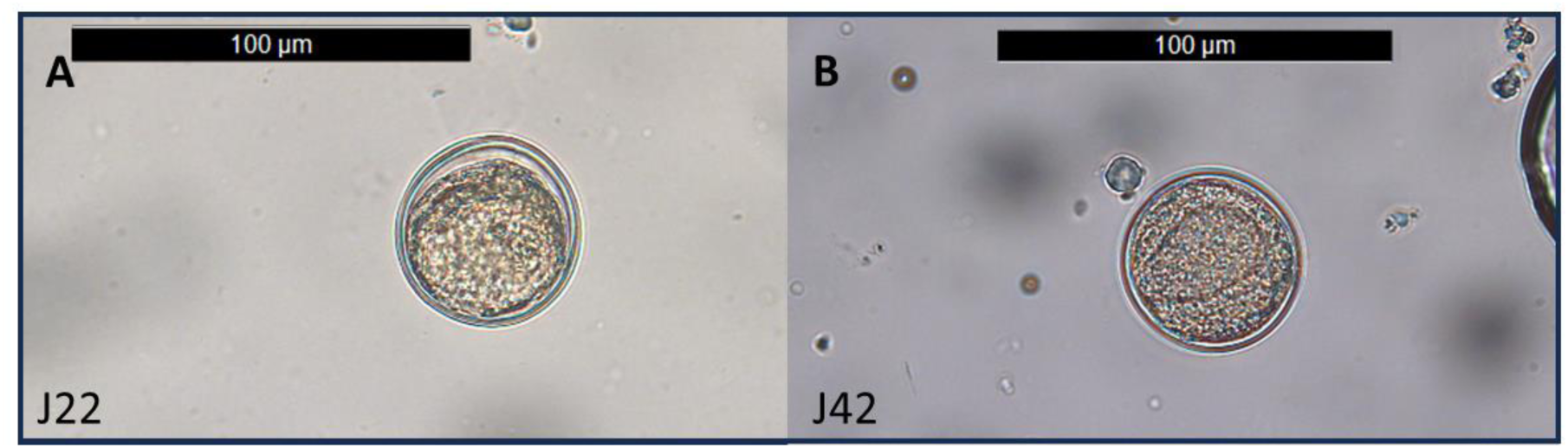
(A) *Anisakis*, as identified through the diagnostic key (Dailey et al. 1980), has a gap between the interior and the outer shell. (B) *Contracaecum* had no gap between the outer wall and the interior. Eggs in (A) and (B) were identified and photographed from a fecal floatation.

### Fecal DNA metabarcoding analysis

Across all 25 fecal samples (*n*_SARKW_ = 3, *n*_NRKW_ = 2, *n*_SRKW_ = 25) that were sequenced, there were a total of 384,109 merged sequence reads after QAQC. Sequencing failure was defined in this case as a sample with <1000 total reads across the entire sample; this occurred in nine samples: four SARKW samples and five SRKW samples (Supplemental Table S1). Further, three samples were collected from the same individual (SRKWs J42 and L86) on the same or consecutive days. It is unlikely for an individual to pass an infection in that time period, so these were considered biological replicates (samples from the same individual within the same 24-hour period). Their information is included in Supplementary Table S1, but only one sample for each individual was used in analysis. When the data were filtered to remove samples with sequence failures or sequences with less than 98% certainty of match to the reference database, the total number of reads was reduced to 335,294 (min = 1,297, max = 43,806, mean = 14,537) across 22 samples. The only confirmed killer whale parasite among the species identified based on metabarcode sequence similarity was *Anisakis* spp. (Figure S1). When subset to only include *Anisakis* spp., there were 312,496 reads (min = 0, max = 43,242, mean = 13,417), or 92.3% of the total reads. *Anisakis* spp. was detected in 17 of the 20 SRKW samples, 1 of the 2 NRKW samples, and 3 of the 3 SARKW samples (Table S1). The most notable off-target detections were for bacteria and fungi species.

Four individuals were sampled on multiple occasions; of these, targeted sequence data were successfully generated (>1000 sequence reads post-QAQC) for two of the individuals (Table S1). For both of these individuals with two successfully sequenced samples, the samples were collected approximately one year apart; one of them was infected on both sampling occasions, and the other was infected on the first sampling occasion, but no parasites were detected on the second sampling occasion.

### Comparing morphological and genetic methodologies

Based on the egg counts obtained from killer whale samples analyzed through sedimentation, 76.5% (26/34) had anisakid eggs present in their fecal samples (Figure 4a). In comparison, *Anisakis* spp. was detected in 94.1% of fecal samples through genetic analysis. Of the six samples that failed to sequence and were removed from the fecal dataset, eggs were not observed in XX. This gives us some indication that sequencing failure may have been partially caused by the absence of target taxa in these samples; however, eggs were observed in XX samples that failed to sequence. Because the primers we used for this study are also known to effectively sequence many species of bacteria and fungi as well as the species of interest, we were not able to distinguish the cause of sequencing failure. Sequencing appeared to be better at detecting *Anisakis* presence than laboratory fecal sedimentation, given the number of samples that had zero eggs but some amount of *Anisakis* DNA detected (Figure 4b). However, low positive read counts should be interpreted with caution as false positive detections can be caused by, e.g., contamination in the lab or by a phenomenon called index hopping, in which sequence reads from one sample sample are assigned to another during the sequencing process.

**Figure 4:**
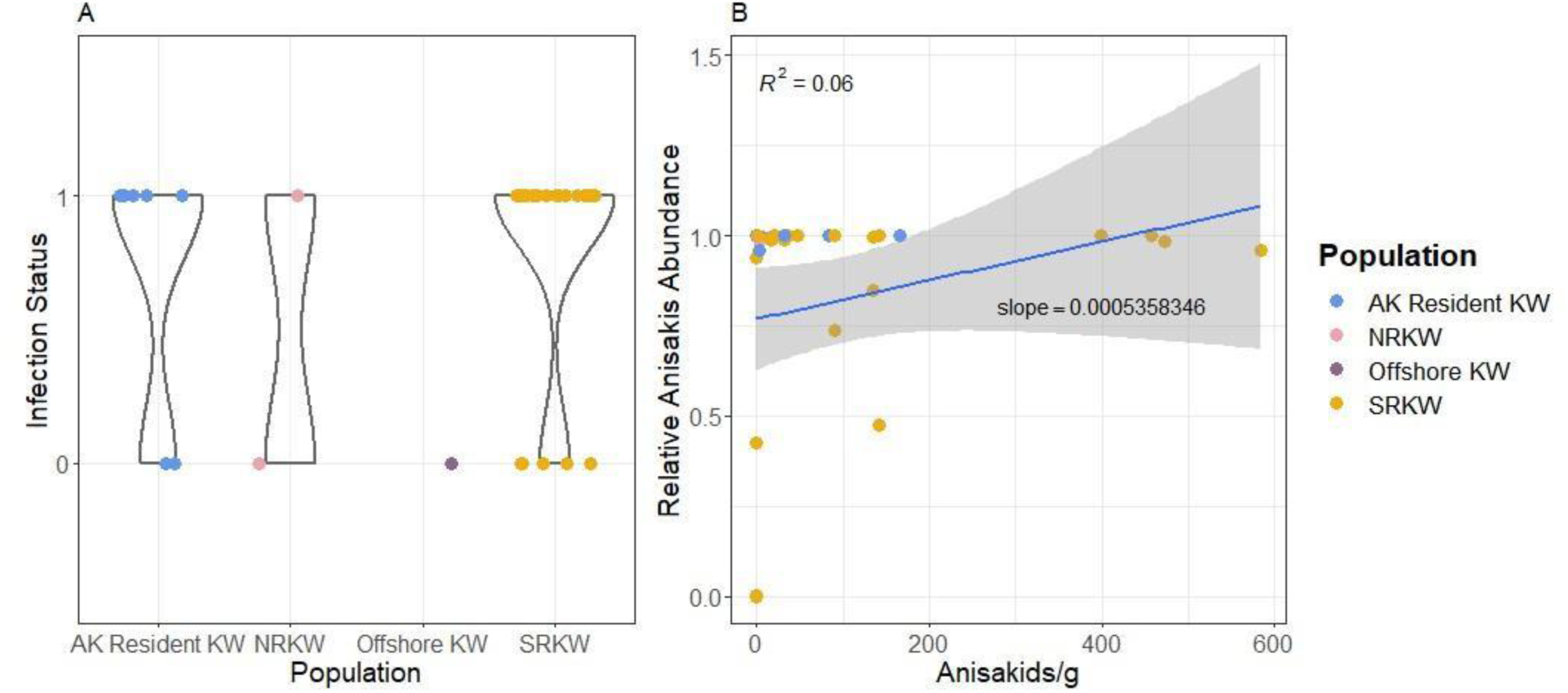
A) The infection status of each population represented by violin plots where 1 indicates that there were anisakid eggs present in the sample (infected), and 0 indicates that anisakid eggs were absent (uninfected). The width of the polygon reflects the number of individuals of a certain infection status for each group. Color indicates the population from which the sample was collected. B) The weak relationship between egg counts of anisakids per gram, and *Anisakis* relative abundance in sequence read counts, determined through eDNA analysis.

### Statistical analysis

#### Between-population differences

The first model assessed whether there was a difference in infection status (i.e., presence of anisakid eggs) among the three resident populations of killer whales sampled. Because anisakids were the only marine mammal parasites occurring across all populations, we only ran the model with the response variable of anisakid infection status. We found that anisakid infection status was not significantly different among populations (Table 2).

**Table 2:** Model 1 assessed differences in anisakid infection status among the resident populations sampled. This analysis used data from all samples with sufficient molecular data available from SARKW (n = 7), NRKW (n = 2), and SRKW (n = 25). Alaska residents were in the reference position.

|  | Estimate | Std. Error | Z value | P value |
| --- | --- | --- | --- | --- |
| Intercept | 0.916 | 0.837 | 1.095 | 0.273 |
| NRKW | -0.916 | 1.643 | -0.558 | 0.577 |
| SRKW | 0.470 | 0.975 | 0.482 | 0.630 |

#### Demographic or temporal effects on likelihood of infection

Our second model assessed whether anisakid infection status was influenced by temporal or demographic variables. Model selection provided the most support for the model that included month as the only fixed effect and whale ID as a random effect (AICc = 28) (Table 3). However, in this model, month did not have a significant effect (estimate = 2.43, SE = 1.376, Z = 1.765, P = 0.078) on the relative abundance of *Anisakis* in a sample.

**Table 3:** Model 2 tested whether demographic variables (age, sex, pod) or temporal variables (month or year) affected anisakid infection status of the host. Model selection suggested a model with month as fixed effect was the best fit. The model was run with November in the reference position, and used all available data from SRKW with known identities and age classes (n = 24).

|  | Estimate | Std. Error | Z value | P value |
| --- | --- | --- | --- | --- |
| Intercept | -0.693 | 1.225 | -0.566 | 0.571 |
| September | 2.428 | 1.376 | 1.765 | 0.078 |

#### Body condition and parasite abundance

Our third model assessed whether SRKW body condition class influenced anisakid infection status (i.e., presence/absence of anisakid eggs). There was no difference in parasite infection status across body condition (Table 4; Figure 4). The model containing body condition was not the best fit model of the candidate models tested (AICc = 23.8, ΔAIC = 2.46), a model without any fixed effects was the best fit (AICc=21.3). The ordinal regression model similarly found no effect of parasite infection status on body condition (estimate = 2.153, SE = 1.617, Z = 1.331, P = 0.183). A power analysis showed that at our current sample size, we would be able to detect a high correlation (0.8) between body condition and parasite infection status 80% of the time. We would have a 17.2% probability of detecting a low correlation (0.3), and a 42.8% chance of detecting a moderate correlation (0.5) (Supplementary Materials).

**Table 4:**
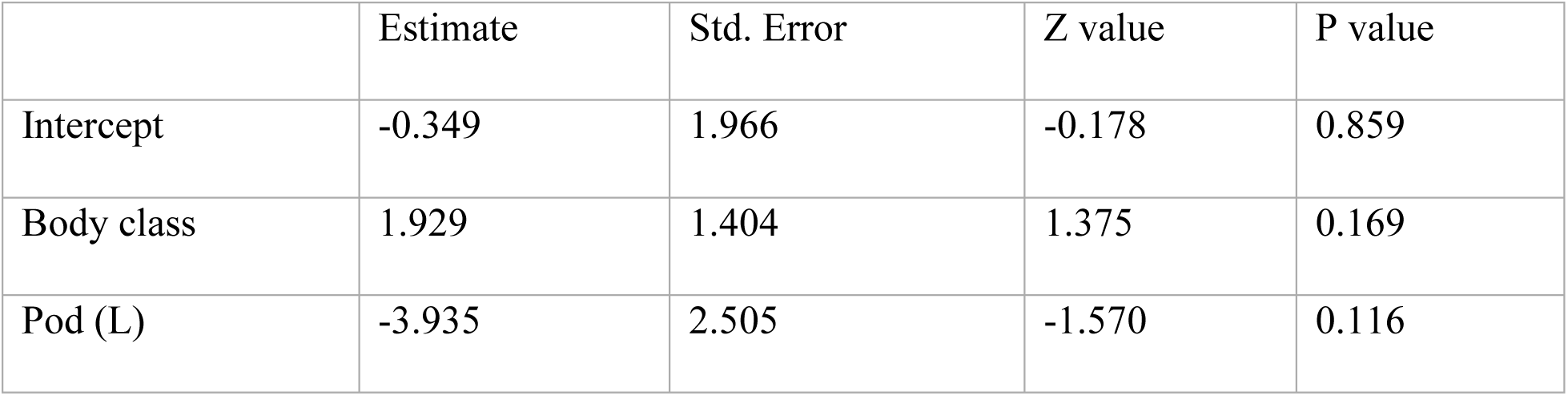
Model 3 assessed whether body condition class affected anisakid infection status. The best fit model was the null, the second-best fit included only Pod (ΔAIC =1.98), and the third included body class and Pod (ΔAIC =2.46). Because we were interested in the effect of body class, we reported the results of the third best fit model. J Pod was in the reference position.

|  | Estimate | Std. Error | Z value | P value |
| --- | --- | --- | --- | --- |
| Intercept | -0.349 | 1.966 | -0.178 | 0.859 |
| Body class | 1.929 | 1.404 | 1.375 | 0.169 |
| Pod (L) | -3.935 | 2.505 | -1.570 | 0.116 |

## Discussion

Prior to this study, the only published documentation of the internal parasite fauna of wild killer whales relied upon data collected during necropsies. Here, we provide quantitative evidence of common parasitic infections in living, healthy killer whales using two independent, complementary analytical approaches. *Anisakis* spp. were detected in most whales sampled, including southern residents, southern Alaska residents, and northern resident killer whales, as well as in the single offshore killer whale sampled. This finding is consistent with what is known about *Anisakis* infections in other odontocetes (Colón-Llavina et al. 2009; Margolis and Dailey 1972), but the widespread infection of killer whales in the Northeast Pacific is a new finding. We demonstrate that both traditional morphological approaches and more modern molecular approaches can be used to detect internal parasite infections of wild killer whales and highlight the benefits and limitations of these two complementary approaches (Supplementary Materials 1).

### Parasites detected

While several parasite species were detected (Supplementary Materials 1), the most prevalent parasite species found in all killer whale populations, and detected by all methodologies, were *Anisakis* spp. There were multiple instances in which the egg count was zero, but *Anisakis* DNA was detected through molecular analysis. This may be due to the variability in egg shedding of the parasites over time (Ugland et al. 2004). An anisakid generally lives in its marine mammal host for 37 to 109 days before maturation, and eggs are shed in the last week of the nematode’s life (Ugland et al. 2004). Individual nematodes can produce between 500,000 and over 1 million eggs, depending on their body length (Simard 1997; Ugland et al. 2004). They shed 85% of their eggs within the first 3 days of spawning, and that rate declines as spawning goes on (Ugland et al. 2004). Depending on when the fecal sample is taken, an infected individual might not have any eggs detected in the fecal sample, though they may host living adult worms, which would be detectable with DNA metabarcoding. Additionally, egg counts cannot reflect the presence of male anisakids or immature females. Because genetic metabarcoding detections are agnostic to the life stage of the parasite, this approach offers the opportunity to detect parasites that would go undetected using traditional fecal float methods. The results of this study highlight the value of leveraging archived fecal samples for novel purposes.

The prevalence of infection through DNA detection was too high to successfully run presence/absence models on metabarcoding detection. While not used in our models, DNA metabarcoding detection is promising as a reliable indicator of infection status or parasite load because this method can detect not only eggs in the fecal sample, but also DNA of adult nematodes still living in the gut tract (Berger and Aubin-Horth 2018). However, several important factors will affect the interpretation of these data and can lead to misleading results if not considered. First, not all parasites known to infect killer whales have sequence data available to detect with molecular methods. In the case of the primers used for this study, sequence failure due to a lack of target species could not be distinguished from sequence failure due to other causes; therefore nine samples were removed from analysis, further limiting the inferences we could make from our small dataset. This is an issue that does not occur in morphological analysis. The relationship between DNA quantity and organismal abundance is unknown (Davey et al. 2021), and observed read counts may be strongly affected by amplification bias (in which some species amplify better than others due to primer mismatch; Shelton et al. 2022) or subsampling bias for rare targets (Pinol et al. 2015; Gold et al. 2023). Additionally, the relative proportion of reads sequenced among various individuals could be biased by the type of genetic material in the sample (worm vs. egg), how accessible DNA content was (i.e., some eggs/stages may be more resilient to lysis), and the composition of the subsample (Gold et al. 2023). While we do not expect our targets to be rare, our samples were subject to many of the remaining challenges so we cannot infer parasite abundance through our metabarcoding analysis. Relative proportion of parasites in a DNA sample can provide some insight into the most common sequences in a sample but are also prone to amplification bias and subsampling bias. Pairing metabarcoding analyses with a qPCR and/or ddPCR methods, as well as the use of mock DNA mixtures with known proportions of input DNA, would facilitate the use of these results to assess the absolute abundance of single species, rather than being constrained to only assessing the relative proportional space of taxa in a mixture. Additionally, continued research comparing DNA metabarcoding with traditional egg counting methods and assessments of gastrointestinal parasite infections in necropsied whales will be important to understanding and interpreting the relationships between DNA metabarcoding data, egg counts, and parasite load. Because both molecular and morphological identification methods are subject to their own biases, employing the two approaches in tandem offers an unique opportunity to leverage the relative strengths of the two methods and further efforts to validate and understand the differences between them.

A low level of contamination was detected in the metabarcoding sequences for some samples. While we did our best to sterilize our sampling tools before sample extraction, we were not working in a sterile environment. Additionally, all fecal samples were collected opportunistically, and collection methods were designed to limit carryover or contamination between samples but were not designed for sterile sample collection or with a focus on parasitic detection. Possible contamination during sample collection cannot be ruled out; however the transparency of off-target contaminating sequences, such as onion and banana, and the rarity of those in the dataset indicates that sample contamination was rare and likely negligible compared to the quantity of DNA from target taxa. Additionally, *Anisakis* does have a free-living stage, from 43–91 days (Measures 1996), before being consumed by an intermediate host or dying; theoretically, some reads could have been attributable to DNA of free-living *Anisakis* in the water.

### Inter-population differences in parasite load

Despite our limited sample size, we expected to detect differences in infection status among the three killer whale populations. Of the resident killer whales in the Northeast Pacific, SARKW and NRKW populations are both larger and increasing, while the SRKW population is small and continues to decline despite targeted recovery actions (Matkin et al. 2014; Fisheries and Oceans Canada 2018). OKWs are more abundant than SRKWs, but little is known about their trends in abundance (Schorr et al. 2022; Ford et al. 2014). Because SRKWs face several cumulative stressors (Lacy et al. 2017; Murray et al. 2021) at levels presumed to exceed that of the other killer whale populations, due to their conservation status, we hypothesized that their immune systems may be compromised (Curry, 1999) compared to the other populations, making SRKW less equipped to resist parasite infections. Contrary to expectations, we did not detect a significant difference in parasite infection status across the killer whale populations we sampled. While our sample size was limited, each population sampled had *Anisakis* spp. infections, with prevalence (i.e., the proportion of infected individuals within a population) of 50% or greater based on molecular identification. This contrasts with the findings of Raverty et al. (2020), in which the authors only detected adult *Anisakis simplex* in one SARKW and 25% (1/4) of the SRKWs necropsied.

The similarity among populations may reflect similarities in the infection prevalence of their prey, and thus their risk of exposure. The three resident populations are all known to consume salmonids along the Northeast Pacific coast (Matkin et al. 2014; Olesiuk et al. 2005; Hanson et al. 2021) and the offshore whales consume higher trophic level fish and sharks (Schorr et al. 2022; Ford et al. 2014). In a recent study, Mastick et al. (2024) found that anisakid abundances in chum and pink salmon have increased in Alaska from 1979 to 2020. As marine mammal populations have increased along the coasts and inland waters of Washington, Oregon, and California (Calambokidis and Baird, 1994; Elliser and Hall, 2021; Jefferson et al. 2021; Calambokidis et al, 2017; NOAA 2017; Brown et al. 2005; Derville et al. 2022; Laake et al. 2018; Lowry, 2014), the prevalence of these nematodes are likely to increase in response. This has been observed in other systems (e.g., Buchmann and Kania 2012; Haarder et al. 2014; Mastick et al. *in revision*). Increased abundances of other species of marine mammal hosts may have contributed to high abundances of anisakids, and therefore to the detection of infection prevalence in the three resident killer whale populations sampled in this study.

#### Demographic or temporal effects on likelihood of infection

Month was the best predictor for anisakid infection status in SRKW, although this parameter was not considered a significant driver of infection status. Given that samples collected in September have an observably higher rate of parasite presence than those collected in November, the non-significance of this result may be due to small sample size; only three samples were collected in November (Figure 5, Table 4). This temporal difference could be due to seasonal differences in SRKW diet (Hanson et al. 2010). From June to August, Chinook salmon is the primary component of SRKW diet. In September, SRKW diet is made up of Chinook and coho salmon (Ford et al. 2016), while in the fall and early winter, their diet is a mixture of Chinook, coho, and chum salmon, as well as non-salmonid species (Hanson et al. 2021; Van Cise et al. 2024).

**Figure 5:**
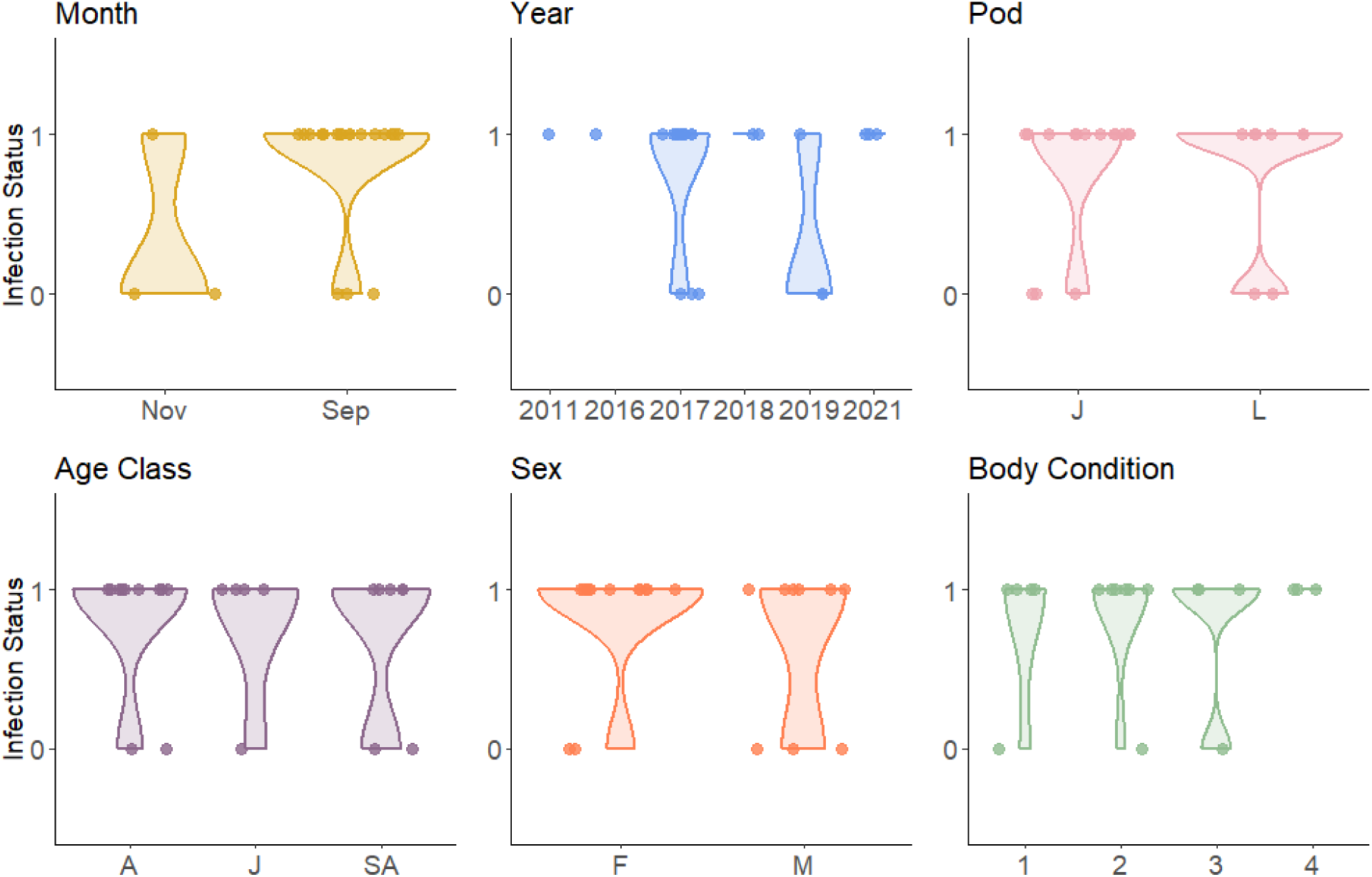
The relationship between anisakid infection status and temporal (year, month) and demographic (pod, age class, sex) factors, as well as body condition class, represented by violin plots. The width of the polygon reflects the number of individuals of a certain infection status for each group. The only factor that had a marginally significant effect on parasite infection status was month. Pod indicates whether the sample was collected from a member of J or L pod (K was not well-represented), and age class indicates if the individual was a juvenile (J), subadult (SA), or adult (A).

Because there is a lag of 37–109 days post-infection for *Anisakis* to reproduce (Ugland et al. 2004), these higher *Anisakis* abundances in the late summer could indicate variable infection levels driven by differences in seasonal diet and species-specific prey infection rate.

#### Anisakids and body condition

Notably, none of the demographic data included in our model were good predictors for parasite infection status. Body condition did not correlate significantly with anisakid infection status in our models, which may be due to the limited sample size available for this pilot study. A similar study in harbor porpoise detected a significant correlation between body condition and parasite infection status using the stomachs of 97 necropsied porpoises to achieve sufficient statistical power (Ten Doeschate et al. 2017). Limited sample size is a common problem with noninvasive samples from rare species in the marine environment (Cossu et al. 2022; Smith and Wang, 2014). We estimate that, if we sampled the entire population at the same level of replication as in our dataset (i.e., 100 samples from all 74 extant whales), we would have greater power to detect even a weak correlation between parasite abundance and body condition 80.8% of the time.

Additional ongoing collection of fecal samples and body condition data will increase the power to detect correlations between parasite infections and body condition, if those correlations exist.

### Implications

Morphological and molecular methodologies proved complementary for detecting gastrointestinal parasites in fecal samples collected from free-swimming cetaceans. Despite demonstrated success, as the first study to employ molecular methods to identify parasites in living killer whales, there may be value in further primer development and validation for future studies to maximize the return of sequence data for the desired taxa. Off-target detections included bacteria and fungi, taxa that are likely to be present at very high concentrations in fecal samples. Refining the design of PCR primers to target parasites and avoid other non-parasitic gut microbes could be beneficial for recovering nematode detections in low abundance. Additionally, we are limited by a lack of data on the relationship between egg counts, DNA read counts, and actual infection burdens. Future studies could aim to study this relationship in a laboratory setting.

Studying parasite infections in living cetaceans provides a valuable avenue that can be leveraged for near-real-time inferences about energetic stresses at the level of the individual, which can potentially be used to inform adaptive management and intervention strategies. Though we did not detect a correlation between body condition and anisakid infection status, it has previously been shown that intestinal parasites contribute to energetic stress (Shanebeck et al. 2022),. The SRKW population has recently been found to exhibit signs of inbreeding depression (Kardos et al. 2023). Many studies have shown that inbreeding leads to higher infection prevalence (Stevens et al. 1997) and susceptibility (Luong et al. 2007), higher parasite intensity and diminished ability to clear an infection (Smallbone et al. 2016), higher infection severity (Arkush et al. 2002), weakened cell-mediated immune response (Reid et al. 2003), and greater prevalence of gastrointestinal parasites (Cassinello et al. 2001). Here, we did not detect a difference in the prevalence of anisakid infections across populations, despite population level differences in inbreeding, however previously demonstrated effects of parasite infection on the extinction risk of inbred populations (McCallum and Dobson 1995; Smallbone et al. 2016) have been noted and warrant consideration in the management of endangered species (Cassinello et al. 2001) and additional research to determine the impact of parasite burden on killer whale fitness.

Anisakid infections are not a permanent ailment, with maximum life spans for *Anisakis* of 109 days within a marine mammal host (Ugland et al. 2004). However, given the increasing abundance of anisakids in salmon (Mastick et al. 2023b) and in the Puget Sound (Mastick et al. *in revision*), as well as the immunocompromised nature of salmon in SRKW foraging region (Sures 2008), these infections may become more prevalent and have a greater impact in the future. *Anisakis* spp. may have a larger energetic cost than previously assumed (Shanebeck et al. 2022) and should be considered in future studies of sublethal stressors in the population.

Parasite infections are rarely monitored in living cetaceans but given the impact of intestinal helminths on hosts (Shanebeck et al. 2022), these data lay the groundwork for monitoring individual health in endangered populations. In the SRKW population, proposed energy deficiencies (Couture et al. 2022; Wasser et al. 2017), high load of pollutants, and inbreeding depression may be impacting immune function in members of this population (Krahn et al. 2009; Kardos et al. 2023). Monitoring the prevalence of gut parasite infections may offer an avenue for detecting the physiological effects of these cumulative stressors. Our study provides valuable new tools for monitoring individual health in living whales and further validation is warranted to understand the importance of parasite identification as another monitoring tool in our management toolbox.

## ACKNOWLEDGEMENTS

Aerial and boat-based operations around whales were conducted under the authority of National Marine Fisheries Service Permits 16163, 19091 and 22306 in US waters. Aerial field operations and data analysis were conducted with funding support from the National Fish and Wildlife Foundation, the National Oceanographic and Atmospheric Administration (NOAA), Shell, Sea World, SR3, and the SeaWorld and Busch Gardens Conservation Fund. The views and conclusions contained in this document are those of the authors and should not be interpreted as representing the opinions of policies of the U.S. Government or the National Fish and Wildlife Foundation and its funding sources. Mention of trade names or commercial products does not constitute their endorsement by the U.S. Government, or the National Fish and Wildlife Foundation or its funding sources. This work was supported by several awards to CLW, including a CAREER Award from the US National Science Foundation Division of Environmental Biology (NSF Grant Number 2141898), a Research Grant from the Cooperative Institute for Climate, Ocean, and Ecosystem Studies (CICOES), a Sloan Research Fellowship from the Alfred P Sloan Foundation, a University of Washington (UW) Innovation Award, and the UW Royalty Research Fund.

