## Supplemental Materials for "Widespread parasite infections in living resident killer whales in the Northeast Pacific Ocean"

### Supplementary Materials

#### Fecal sample collection

Fecal samples from wild killer whales have been collected opportunistically as part of long-term monitoring of the Northeast Pacific killer whale RKW populations killer whale populations. Fecal samples were collected from all populations using methodology previously described by Hanson et al. (2010; 2021) and Ford et al. (2016). Molecular genetic analyses were performed on each fecal sample to identify the host whale, as described in Ford et al. (2016). SRKW samples were assigned to individual whale identities using a custom killer whale nuclear single nucleotide polymorphism (SNP) panel to generate multilocus genotypes for each individual fecal sample and assign whale identity with reference to genotypes generated from vouchered (i.e., known whale identity) tissue biopsies. The genetic reference database for SRKWs includes approximately 90% of extant individuals in the population. As such, each SRKW fecal sample is linked to an individually identified whale. SARKW and NRKW samples were collected in separate field efforts and were not identified to the individual level.

#### Fecal float and sedimentation analysis

Briefly, each sample was reconstituted with a soap solution (30 ml of Dawn soap in 1 gallon tap water) and mixed until thoroughly emulsified. The solution was strained using gauze into a 50 mL conical tube and centrifuged at 1300 rpm for 10 minutes in a fixed-head centrifuge. Several drops of the resulting pellet were subsampled for sedimentation. We then decanted the soap solution from the pellet and reconstituted the pellet with zinc sulfate solution (1.18 specific gravity). The mixture was transferred to a 15-mL conical tube and filled to 14.5 mL, then centrifuged for 1300 rpm for 10 minutes. The tube was carefully removed from the centrifuge and filled until there was a slight positive meniscus. We placed a coverslip on top of the tube and waited 10 minutes before removing the coverslip for immediate analysis. Both sedimentation and floatation slides were examined under a Leica DM2500 compound microscope to count and photograph parasite eggs.

**Fecal DNA metabarcoding analysis**

We ran an independent molecular genetic approach to detect parasites and pathogens. We sent the remaining portion of each fecal sample, which varied depending on the starting weight, to Jonah Ventures environmental genetics lab in Boulder, Colorado, for library preparation and sequencing. Frozen samples were thawed for 1 to 2 hours before subsampling for DNA extraction. Under a laminar flow hood, sterile cotton swabs (Fisher, cat# 22-363-173) were coated with fecal matter, and the swabs were placed in labeled extraction tubes. Sterile tweezers and pliers were used to handle cotton swabs and remove the wooden ends of the cotton swab before extraction. Samples were immediately processed or stored at –20°C until extraction.

Genomic DNA was extracted from samples using the DNeasy 96 PowerSoil Pro Kit (384) (Cat # 47017) according to the manufacturer’s protocol. Genomic DNA was eluted in 100 µl of elution buffer and frozen at –20°C until the DNA amplification step. We amplified a 156 bp fragment of the 28S region of ribosomal RNA using a published primer set that targets eukaryotic organisms including Apicomplexa, Phragmoplastophyta, Ascomycota, Basidiomycota, Annelida, Arthropoda, Mollusca, Nematoda, and Platyhelminthes: nucLSUDf1 (5'-CGTCTTGAAACACGGACCAAG-3') and nucLSUDr1 (5'-GCATAGTTCACCATCTTTCGGG-3'), to amplify the forward and reverse segments, respectively (Sonnenberg et al. 2007; Cabodevilla et al. 2022). This primer was selected based on its likelihood of amplifying all but one of the parasite species likely to be found in these killer whale populations (*Campula* spp., for which no reference sequence was available; Table 1), and ability to differentiate to the genus or species level for all of the possible parasites. This primer includes all other known parasite species (Cabodevilla et al. 2022). However, this primer has been shown to perform poorly at detecting Phragmoplastophyta, Arthropoda, Nematoda, Platyhelminthes, and Apicomplexa, e.g. only detecting 33% of Platyhelminths species tested (Cabodevilla et al. 2022). Both forward and reverse primers were modified to include a 5' adaptor sequence to allow for subsequent indexing and Illumina sequencing.

Target amplicons were amplified in 25-µL PCR reactions according to the Promega PCR Master Mix specifications (Promega catalog # M5133, Madison, WI) which included 12.5 µL Master Mix, 0.5 µL of each primer, 1.0 µL of gDNA, and 10.5 µL DNase/RNase-free H2O. Fecal genomic DNA was amplified using the following conditions: initial denaturation at 94°C for 3 minutes, followed by 35 cycles of 30 seconds at 94°C, 30 seconds at 57°C, 1 minute at 72°C, with a permanent hold at 4°C. To determine amplicon size and PCR efficiency, each reaction was visually inspected using a 2% agarose gel with 5 µl of each PCR as input.

Amplicons were then cleaned by incubation with Exo1/SAP for 30 minutes at 37°C followed by inactivation at 95°C for 5 minutes, and stored at –20°C before the indexing PCR reaction.

A second round of PCR was performed to complete the sequencing library construct, appending with the final Illumina sequencing adapters and integrating a sample-specific, 12-nucleotide index sequence. The indexing PCR included Promega Master mix, 0.5 µM of each primer and 2 µLof template DNA. The indexing cycling protocol consisted of an initial denaturation of 95 °C for 3 minutes followed by 8 cycles of 95°C for 30 sec, 55°C for 30 seconds, and 72°C for 30 seconds. Final indexed amplicons from each sample were cleaned and normalized using SequalPrep Normalization Plates (Life Technologies, Carlsbad, CA). Samples were pooled to equal volume and sequenced on an Illumina MiSeq (San Diego, CA) at the Texas A&M Agrilife Genomics and Bioinformatics Sequencing Core facility using a v2 500-cycle kit (cat# MS-102-2003). Necessary quality control measures performed at the sequencing center prior to sequencing included assessing the size distribution of the pooled sequences and diluting the final sample to a concentration of XXX before loading onto the sequencer.

Initial quality control of the sequencing data, and taxonomic classification of sequences, was completed by Jonah Ventures. Raw sequence data were demultiplexed using *pheniqs* v2.1.0 (Galanti et al. 2021), enforcing strict matching of sample barcode indices (i.e, no errors). *Cutadapt* v3.4 (Martin 2011) was then used to remove PCR primers from the forward and reverse reads, discarding any read pairs where one or both primers were not found at the expected location (5') with an error rate < 0.15. Read pairs were then merged using *vsearch* v2.15.2 (Torbjorn et al. 2016), discarding resulting sequences with a length of < 300 bp, > 450 bp, or with a maximum expected error rate > 0.5 bp (Edgar and Flyvbjerg 2015). For each sample, reads were clustered using the *unoise3* denoising algorithm (Edgar 2016) as implemented in *vsearch*, using an alpha value of 5 and discarding unique raw sequences observed fewer than 8 times. Counts of the resulting exact sequence variants (ESVs) were then compiled and putative chimeras were removed using the *uchime3* algorithm, as implemented in *vsearch*. For each final ESV, a consensus taxonomy was assigned using a custom best-hits algorithm built by Jonah Ventures and the SILVA reference database (GenBank, Benson et al. 2005, Quast et al. 2013) as well as Jonah Ventures voucher sequences records. The reference database was restricted to exclude killer whale DNA. Reference database searching used an exhaustive semi-global pairwise alignment with vsearch, and match quality was quantified using a custom, query-centric approach, where the percent match ignores terminal gaps in the target sequence, but not the query sequence. The consensus taxonomy was then generated using either all 100% matching reference sequences or all reference sequences within 1% of the top match, accepting the reference taxonomy for any taxonomic level with > 90% agreement across the top hits.

We processed and analyzed the classified read count data received from Jonah Ventures using the *phyloseq* package in R (McMurdie and Holmes 2013). We first filtered out all taxonomic classifications that were classified by the Jonah Ventures algorithm with less than 98% confidence, and any taxa that did not make up at least 1% of one or more samples. We considered read counts below 1000 to be a sequencing failure. Once initial quality filtering was completed, we grouped parasite sequences by genus, and estimated the relative proportion of each genus in a sample as the genus-specific read count divided by the total read count for all sequences in the sample.

In addition to the Anisakid spp. detections reported in the manuscript, we detected two other potential eukaryotic pathogens, including the fungi *Aspergillus penicillioides* and *Candida* spp. Other species in this family cause fungal infections in odontocetes (Gaydos et al. 2004), and the species documented here can cause respiratory disease in humans (Klich, 2009), but it is not known to cause the same pathology in odontocetes (Joe Gaydos, marine mammal veterinarian, pers. comm.). The sequence was only detected in two whales at a low level (i.e., less than 50% relative abundance) (Figure S2). *Candida* spp. was the other genus detected in two different whales at similarly low levels (Figure S2). The fungi *Candida* spp. have caused fungal infections in odontocetes including killer whales, though we could not identify the sequence to the species level, so we cannot be sure that this specific species causes similar pathology in killer whales (Gaydos et al. 2004). Several ciliates and copepods were detected, but not the species that have been associated with skin disease in killer whales (Schulman and Lipscomb, 1999; Vecchione et al. 2014).

In addition to detecting species that could be pathogenic to killer whales, we also detected parasites known to infect salmon. *Henneguya* spp., a myxozoan parasite known to cause disease in salmon (Fiala et al. 2015; Fish, 1939), but not odontocetes (Joe Gaydos, marine mammal veterinarian, pers. comm.) was detected in six samples from three individuals. This suggests that some of the salmon eaten by killer whales were infected with parasites that could be detected through the whale's feces. Additionally, the algae *Heterosigma akashiwo* was detected in one sample at a relatively low level (Figure S2). *H. akashiwo* causes red tides that can be lethal to fish (Khan et al. 1997; Taylor and Haigh 1993), and occurs in areas of the SRKW range (Hard et al. 2000; O'Halloran et al. 2006). It is possible, but unlikely that this detection came from DNA in the water surrounding the sample when collected.

No *Contracaecum* spp. were detected in the genetic analysis, though the selected primer should have been able to differentiate the two anisakid species. Though this could have occurred if *Contracaecum* spp. were rare enough that they were not sequenced or due to poor egg lysis during DNA extraction, the life history of *Contracaecum* spp. provides support for this being a misidentification. *Contracaecum* spp. are not known to use cetaceans as definitive hosts and are not expected to reach a reproductive adult stage within a killer whale (Klimpel and Palm 2011), therefore it would be unlikely to detect eggs in killer whale feces. This leads us to hypothesize that following the diagnostic key may lead to misidentification of eggs in killer whale fecal samples.

We also detected a few sequences that suggested there may have been contamination following collection of the fecal samples. For example, in one sample we detected onion sequences (*Allium cepa*, relative proportion in one sample = 0.003), and in another we found banana (*Musa acuminata*, relative proportion in sample = 0.004). These may have come from contamination upon collection, storage in the field, or during sub-sampling and analysis.

**Power Analysis**

We performed a power analysis for Model 3 to quantify the statistical power for detecting effects of body condition on parasite infection status in two simulated datasets: one with our sample size (19 fecal samples from 14 whales), and one that sampled every extant individual with the same proportion of repeated individuals as was observed in our dataset (100 fecal samples from 74 whales). Because whales were sampled more than once, we generated our simulated data based on the same ratio of unique whales to fecal samples (14/19 = 74/100). We simulated our datasets based on the six years when both photogrammetry and fecal data were available. To inform the simulation, we used existing demographic data from the Orca Network. We created a dataset of the IDs, associated pods, and birth years of each individual in the population alive from 2008 to

2021. For each row, we assigned a random year from one of six, representing the six years in which our samples were obtained. We subtracted the birth year from the assigned year to get age at sampling and reassigned any negative values to 0. We assigned age classes to each individual, including 1 (calves, 0–4 years of age), 2 (subadults, 5–11 years of age), 3 (adults, 12+ years of age). Age class was treated as a factor. We simulated body condition data based on the true measurements in our dataset for each age class (i.e., all members of the same age class are assigned a body condition sampled from a distribution of the frequency of body conditions observed in our dataset for individuals in that group), and assigned measurements based on age class. We simulated a parasite presence/absence vector using a binomial distribution based on the presence/absence ratio from our data.

Using the simulated data, we built a simplified GLMM with centered and scaled infection status as the response variable, scaled body condition as a fixed effect, and whale ID as a random effect. Both variables were scaled using the scale() function in base R. By centering and scaling the values of the predictor and response variables, we standardized the regression so that the beta coefficients each represent a correlation coefficient. We ran the model with 500 bootstrap iterations to estimate the likelihood of obtaining a value of $p < 0.05$ using the simr package (Green and MacLeod 2016). We tested a range of potential correlation coefficients between body condition and infection status, from 0.1 (very small correlation) to 0.9 (very large correlation) (i.e., Mukaka 2012) by manually changing the fixed effect coefficient in the model and running the simulation to determine how much power we had to detect an effect. We first ran this analysis with our simulated small dataset (19 fecal samples from 14 whales) and then repeated the analysis with our simulated large dataset (100 fecal samples from 74 whales). We ran the model through the powerCurve() function in the simr package (Green and MacLeod 2016) to determine the sample size needed to detect  low (0.3), moderate (0.5), or high (0.7) correlation between body condition and infection status on the large dataset.

The power analysis demonstrated that at our current sample size, we would be able to detect a high correlation (0.8) between body condition and relative parasite abundance 80% of the time. If there was a low correlation (0.3), we would have only a 17.2% probability of detecting it at our current sample size. We would have a 42.8% chance of detecting a moderate correlation (0.5; Table S2). If we were to sample every whale in the population with the same level of replication as observed in our data (n = 100), we would be able to detect even a low (0.3) correlation between relative parasite abundance and body condition 80.8% of the time.

**Non-anisakids detected**

Several parasites were detected morphologically that were not identified by molecular methods. Through fecal floatation, we detected what was likely a trematode of the genus *Odhneriella* from the OKW sampled (Figure S3a). Through fecal sedimentation, we found cysts consistent with *Balantidium* spp. in SRKW samples (Figure S3b). *Balantidium* spp. was not detected genetically, though it should have been amplified by the selected primer. *Balantidium* spp. are protozoan parasites that have previously been detected in fin whales in Portugal (*Balaenoptera physalus*; Hermosilla et al. 2016), but to our knowledge this group has not been described in delphinids in the Northeast Pacific. The only species of this family known to be pathogenic for mammals is *Balantidium coli* (Ponce-Gordo et al. 2011), which commonly infects terrestrial animals (Schuster and Ramirez-Avila 2008), and has only been detected in one marine mammal species, the Chilean sea lion (*Otaria flavescens*; Hermosilla et al. 2013). We could not definitively identify the cysts using morphological keys, so it is unclear whether the cysts identified in our analysis are *B. coli* and thus, potentially pathogenic. While *Odheriella* spp. was included in our list of possible parasites (Table 1), a genetic sequence was not available on GenBank to use for reference in molecular identification. From our molecular analysis, we did not detect any additional parasite species of whales (Table 1).

We detected several eggs through sedimentation and floatation that we were unable to identify through morphological identification (Supplementary Materials 2: Figure S2). As these eggs were rare and microscopic (<50μm), we were unable to isolate and sequence them independently. Our inability to identify these eggs is probably due to the limited scope of the marine mammal parasite taxonomic identification resources for parasite eggs of delphinids in this region. To continue to improve our ability to diagnose cetaceans through fecal samples, additional efforts are needed to improve morphological identification of fecal parasites, sequence unknown taxa and analyze their phylogenetic and phylogeographic positions, and work to develop primers that capture all known parasites.

**Supplementary Tables:**

Table S1: The anisakid prevalence in each sample, from sedimentation (Anisakids/g), sequencing (*Anisakis* read counts), and the relative proportion of *Anisakis* reads to other identified sequences in each sample. Colors indicate individuals that were sampled more than once.

| Whale ID | Population | Pod | Date | Anisakids/g | *Anisakis read counts* | Relative Proportion of Anisakis reads | Total reads after QAQC filtering |
|---|---|---|---|---|---|---|---|
| ARKW_12 | ARKW | UNK | 2017 | 82.9 | 37530 | 0.999 | 37579 |
| ARKW_13 | ARKW | UNK | 2017 | 31.4 | 18 | 1 | 18 |
| ARKW_15 | ARKW | UNK | 2017 | 165.9 | 38844 | 1 | 38844 |
| ARKW_28 | ARKW | UNK | 2018 | 0 | 0 | NA | 0 |
| ARKW_27 | ARKW | UNK | 2018 | 6.5 | 43242 | 0.988 | 43806 |
| ARKW_03 | ARKW | UNK | 2018 | 2.3 | 451 | 0.982 | 524 |
| ARKW_01 | ARKW | UNK | 2018 | 0 | 56 | 1 | 73 |
| J26 | SRKW | J | 28 Sep 2011 | 20.2 | 142 | 1 | 142 |
| K25 | SRKW | K | 30 Oct 2013 | 134.9 | 8312 | 0.996 | 8342 |
| UNK | SRKW | U | 22 Sep 2015 | 2 | 49 | 1 | 49 |
| J42 | SRKW | J | 14 Sep 2016 | 90.7 | 25576 | 0.999 | 25596 |
| L106 | SRKW | L | 10 Sep 2017 | 0 | 3847 | 0.422 | 9145 |
| J35 | SRKW | J | 19 Sep 2017 | 36 | 7980 | 0.999 | 8009 |
| L113 | SRKW | L | 22 Sep 2017 | 48.2 | 2341 | 1 | 2341 |
| L47 | SRKW | L | 23 Sep 2017 | 0 | 12 | 0.002 | 6621 |
| J42 | SRKW | J | 23 Sep 2017 | 0 | 3378 | 0.938 | 3600 |
| J42 | SRKW | J | 23 Sep 2017 | 91.2 | 4480 | 0.734 | 8321 |

| | | | | | | | |
|---|---|---|---|---|---|---|---|
| J42 | SRKW | J | 24 Sep 2017 | 2506.5 | 18710 | 1 | 18710 |
| J36 | SRKW | J | 24 Sep 2017 | 472.5 | 10932 | 0.982 | 11127 |
| J22 | SRKW | J | 24 Sep 2017 | 584.9 | 3647 | 0.948 | 3923 |
| L86 | SRKW | L | 26 Sep 2017 | 134.9 | 13162 | 0.848 | 15626 |
| L86 | SRKW | L | 26 Sep 2017 | 458.1 | 36287 | 0.999 | 36333 |
| J19 | SRKW | J | 26 Sep 2017 | 14.8 | 7122 | 0.991 | 7223 |
| L118 | SRKW | L | 30 Sep 2017 | 32.2 | 10618 | 0.987 | 10758 |
| J16 | SRKW | J | 13 Sep 2018 | 399.6 | 38972 | 1 | 38982 |
| J49 | SRKW | J | 25 Sep 2018 | 135 | 1387 | 0.814 | 1724 |
| UNK | NRKW | UNK | 19 Aug 2019 | 2.7 | 9917 | 0.989 | 10049 |
| UNK | NRKW | UNK | 19 Aug 2019 | 0 | 0 | 0 | 1297 |
| J49 | SRKW | J | 13 Nov 2019 | 0 | 0 | 0 | 7008 |
| J39 | SRKW | J | 13 Nov 2019 | 141.4 | 45 | 1 | 45 |
| J27 | SRKW | J | 13 Nov 2019 | 0 | 0 | 0 | 18 |
| L116 | SRKW | L | 11 Sep 2021 | 21.3 | 3553 | 0.979 | 3650 |
| L106 | SRKW | L | 12 Sep 2021 | 141.2 | 53 | 0.53 | 112 |
| J26 | SRKW | J | 15 Sep 2021 | 17.2 | 17347 | 0.983 | 17685 |

Table S2: The estimated power to detect varying correlations ranging from 0.1 to 0.9 of anisakid infection status with body condition. We ran a power analysis for our sample size ($N_{fecals}$ = 19) and for a scenario in which all of the whales in the population were sampled with the same replication ($N_{fecals}$ = 100). Power estimates were calculated with the powerSim() function in the simr package in R.

| Correlation coefficient | Power ($N_{fecals}$ = 19) | Power ($N_{fecals}$ = 100) |
|---|---|---|
| 0.1 | 4.4% | 17.0% |
| 0.3 | 17.2% | 85.2% |
| 0.5 | 42.8% | 100% |
| 0.7 | 72.8% | 100% |
| 0.9 | 89.8% | 100% |

Supplementary Figures:

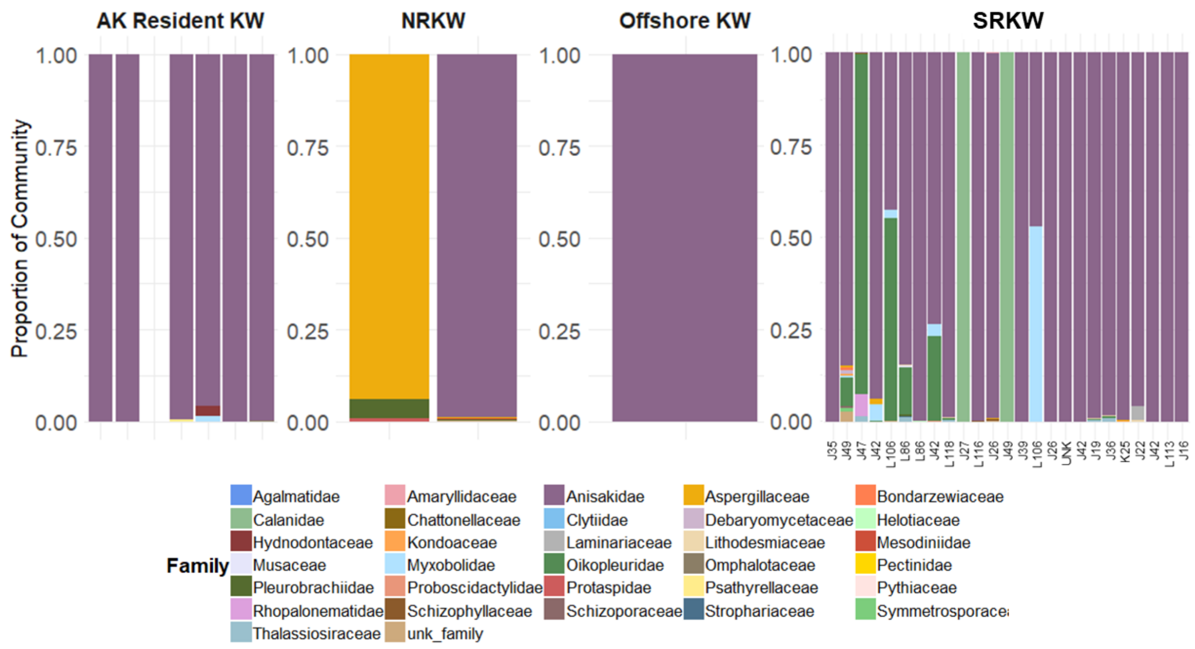


Figure S1: The relative abundance of sequences by family detected in each sample varied by population. Each bar represents an individual sample. Anisakidae was the most commonly detected sequence in our molecular analysis. Unk_family indicates a sequence that we could not identify at the family level. Besides Anisakidae, none of the other families are known to be pathogenic to killer whales.

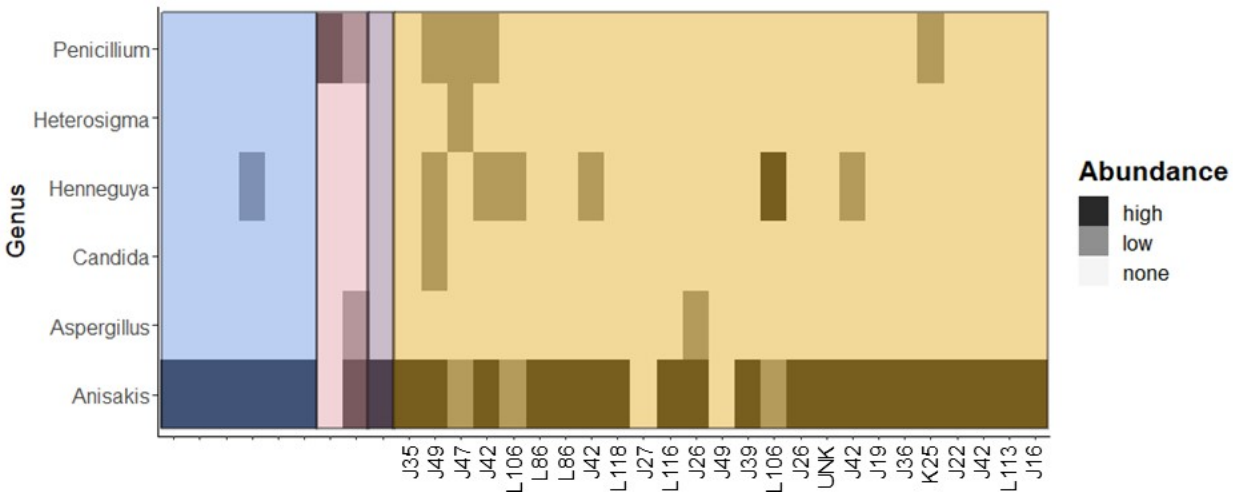


Figure S2: A heatmap of the relative abundance of each potentially pathogenic genus detected across all samples using metabarcoding. Blue indicates SARKW, pink is NRKW, purple is OKW, and yellow is SRKW. High relative abundance indicates read counts for a given genus make up 50% or more of the total sample, while low relative abundance indicates the read counts made up less than 50% of the samples.

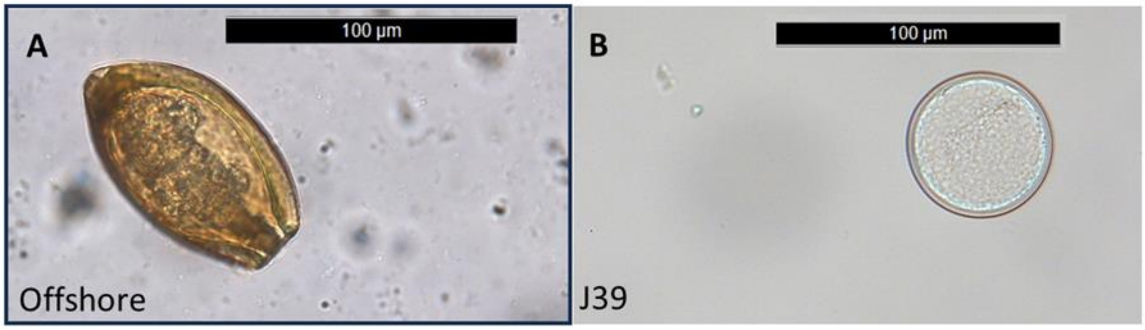


Figure S3: (C) Likely *Odhneriella* spp., found in the offshore killer whale sample. (D) Possibly *Balantidium* spp., which has previously been found in fin whales (Hermosilla et al. 2016). Eggs in (C) were identified and photographed from a fecal floatation, egg in (D) was identified and photographed from a fecal sedimentation.

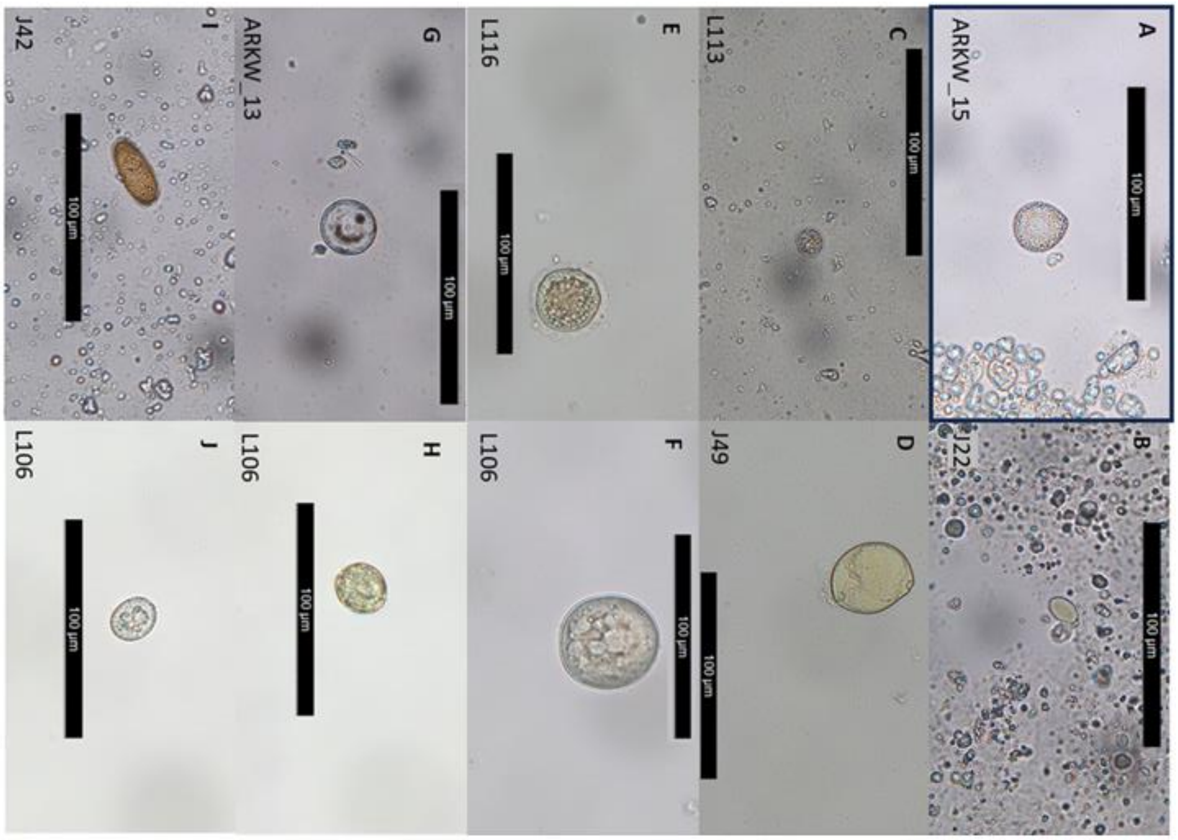


| | Total eggs found in the population's samples | | | |
|---|---|---|---|---|
| **Parasite** | **ARKW** | **NRKW** | **Offshore** | **SRKW** |
| Unknown A | 2 | 0 | 0 | 0 |
| Unknown B | 0 | 0 | 0 | 2 |
| Unknown C | 7 | 0 | 0 | 13 |
| Unknown D | 0 | 0 | 0 | 1 |
| Unknown E | 0 | 0 | 0 | 2 |
| Unknown F | 0 | 0 | 0 | 1 |
| Unknown G | 1 | 0 | 0 | 0 |
| Unknown H | 0 | 0 | 0 | 1 |
| Unknown I | 0 | 0 | 0 | 1 |
| Unknown J | 0 | 0 | 0 | 1 |

Figure S3: Unidentifiable parasites were found in fecal sedimentation and floatation in small numbers.
